# Determinants of precocious B-cell aging in European adolescents living with perinatally acquired HIV-1 after over 10 years of suppressive therapy

**DOI:** 10.1101/2021.11.11.468189

**Authors:** Alessandra Ruggiero, Giuseppe Rubens Pascucci, Nicola Cotugno, Sara Domínguez-Rodríguez, Stefano Rinaldi, Alfredo Tagarro, Pablo Rojo Conejo, Caroline Foster, Alasdair Bamford, Anita De Rossi, Eleni Nastouli, Nigel Klein, Elena Morrocchi, Benoit Fatou, Kinga K. Smolen, Al Ozonoff, Katherine Luzuriaga, Hanno Steen, Carlo Giaquinto, Philip Goulder, Paolo Rossi, Ofer Levy, Savita Pawha, Paolo Palma, on behalf of the EPIICAL consortium.

## Abstract

HIV infection results in a state of chronic immune activation leading to premature immune aging, B-cells dysfunction, that persists despite prolonged virological suppression. In this scenario, adolescence living with perinatally acquired HIV (PHIV), deserve a peculiar attention since potentially exposed for their entire life to chronic immune activation. Here we identified determinants of precocious aging B cells in 40 PHIV undergoing suppressive antiretroviral therapy (ART) for median 13.5 years. All individuals started ART by 2^nd^ year of life and achieved virus suppression within the 1^st^ year of ART, with majority of patient maintaining suppression until analysis and 5/40 experiencing viral Spike (transient elevation of HIV-1 VL, 50-999 copies/ml). We employed a multiomics approach including deep immunological B and T cell phenotype in PBMC, with aging B cells defined by the expression of T-bet and CD11c; plasma proteomics analysis by mass spectrometry and serum level of anti-measles antibodies as correlates of humoral response. We found that individuals with expansion of aging B cell, defined by the expression of T-bet+CD11c+, were those starting treatment later, presenting detectable levels of cell-associated HIV-1 RNA, history of Spikes, and a higher frequency of exhausted T-cells, including those expressing PD-1, LAG3, TIGIT. Accordingly, the proteomic analysis revealed that subjects with expansion of aging B cells and exhausted T cells had enrichment of proteins involved in immune inflammation and complement activation pathways, such as CLU and APCS which are also involved in tumor progression. Signs of precocious aging were associated with a reduced capacity to maintain virological memory against measles vaccination. To our knowledge, this is the first study focusing on precocious B-cell aging and dysfunctionality in PHIV with long-term virological suppression. Our experimental strategy enabled identification of clinical, viral, cellular and plasma soluble markers associated with B-cells aging. Our results pave the way to further define risk of disease progression or lymphoproliferative disorders in PHIV.

**Author summary:** Despite a successful antiretroviral therapy (ART), adolescence living with perinatally acquired HIV (PHIV) experience B-cells dysfunction, including loss of vaccine-induced immunological memory and higher risk of developing B-cells associated tumors. It is thus paramount to define novel and precise correlates of precious aging B cell for the definition of novel therapeutic strategies. Here, we studied 40 PHIV who started treatment by 2^nd^ year of life and maintain virological suppression for 13.5 years, with 5/40 patients experiencing transient elevation of the HIV-1 load in the plasma (Spike). We applied a multi-omics approach including immunological B and T cell phenotype, plasma proteomics analysis and serum level of anti-measles antibodies as functional correlates of vaccine-induced immunity. We found that levels of aging B cells were positively associated with age at ART start, cell associated HIV-1 RNA (caHIV-1 RNA) and the presence of Spikes. Individuals with increased proportions of aging B cells had concomitant expansion of exhausted T cells and were unable to maintain vaccine-induced immunity over time. B-cell aging, and T-cell exhaustion were also associated with proteins involved in immune inflammation. The factors found here to be associated with aging B-cell could inform further therapeutic studies.

## INTRODUCTION

HIV-1 replication is associated with abnormalities in all major lymphocyte populations, including the B-cell compartment which results in hyperactivation and exhaustion (1-5). While early antiretroviral therapy (ART)-initiation partially averts this detrimental condition (6), late ART initiation during the chronic stage of HIV infection results in precocious aging of the immune system with irreversible loss of memory B cells and expansion of exhausted B cell subsets including activated memory (AM), double negative (DN)- and tissue-like memory B cells (TLM)(1, 4, 6, 7). The adhesion molecule CD11c and the transcription factor T-bet identify a discrete B cell subset, induced by innate activation and maintained by chronic inflammation or antigen stimulation, may play a detrimental role in chronic HIV infection (8). Overall, chronic B cell activation observed during HIV infection has been related to a reduction of functional resting memory B cells resulting in precocious waning of routine vaccine-induced antibody titers (9-11) and increased risk of age-associated pathologies (12, 13), including malignancies (14). Indeed, a B cell lymphoproliferative disorder such as Hodgkin’s Lymphoma has remained stable or even increased in HIV-positive adults since the introduction of ART and is ∼11-fold higher than in the HIV-negative population (15). In this context, perinatally HIV infected children deserve particular attention, given their life-long exposure to chronic immune activation. It remains unknown whether early ART initiation during acute HIV infection followed by long-term virological suppression could prevent precocious aging of the B-cell compartment. Longitudinally well characterized, adolescents living with perinatally acquired HIV-1 (PHIV) with sustained and prolonged virological suppression represent a unique opportunity to investigate this scientific question. Indeed, children who started ART in infancy are rarely able to achieve and consistently maintain viral control for long periods. In the present work, we attempt to identify determinants of B-cell activation and dysfunctionality in European PHIV who have been treated with ART for >13 years and have a documented history of virus suppression. We performed deep B and T cell phenotyping with a particular focus on factors associated with lymphocyte aging and extensive mass spectrometry-based plasma proteomic analysis. Serum levels of anti-measles antibodies (Abs) were analyzed as correlates of functional humoral immune response.

## RESULTS

### Study cohort

Patient characteristics are shown in Table 1. Overall, we analyzed 40 PHIV (males 13/40, 32.5%), that started ART at a median 4.1 months (IQR 0.3-6.2), achieved virological suppression after median 4.69 (2.52–6.26) and were successfully on ART for median 13.5 years (8.1-16.5). We measured cell associated (ca)HIV-1 DNA (caHIV-1 DNA median 48.8 copies/10^6^ PBMC), caHIV-1 RNA in the Pol and LTR regions. Overall, 5/40 (13%) had experienced a Spike in HIV-1 Viral Load (HIV-1 VL between 400-999 c/mL, returning to VL <50 c/ml at next blood draw) at some point in their lives (Table 1, Fig.1).

**Table 1.** Characteristics of the study population.

|  | CARMA COHORT |
| --- | --- |
| N | 40 |
| Gender M (%) | 13/40 (32.5%) |
| At ART start |  |
| Age median months (IQR) | 4.1 (0.3-6.2) |
| CD4<sup>+</sup> T cells percentage (IQR) | 30.5 (19.2-42.5) |
| Plasma HIV-1 RNA median copies/μL (IQR) | 5.3 (4.1-5.7) |
| Time to suppression median months (IQR) | 4.69 (2.52–6.26) |
| At analysis |  |
| Age (years) | 13.5 (8.7-16.6) |
| Time on ART (years) | 13.5 (8.1-16.5) |
| CD4<sup>+</sup> T cells percentage (IQR) | 41.0 (33.8-46.2) |
| Spike yes or no, n (%) | 5/40 (13%) |
| caHIV-1 DNA (copies/10<sup>6</sup> PBMCs) | 48.4 (6.7-112.5) |
| caHIV-1 RNA (Pol) (copies/10<sup>6</sup> PBMCs) | 0.0 (0.0-1.4) |
| caHIV-1 RNA (LTR) (copies/10<sup>6</sup> PBMCs) | 2.7 (0.0-44.1) |
| anti-Measles IgG, median IU/l (IQR) | 617 (411-936) |
| anti-Measles IgG, median years from vaccination (IQR) | 5 (2-8) |

**Fig.1.**
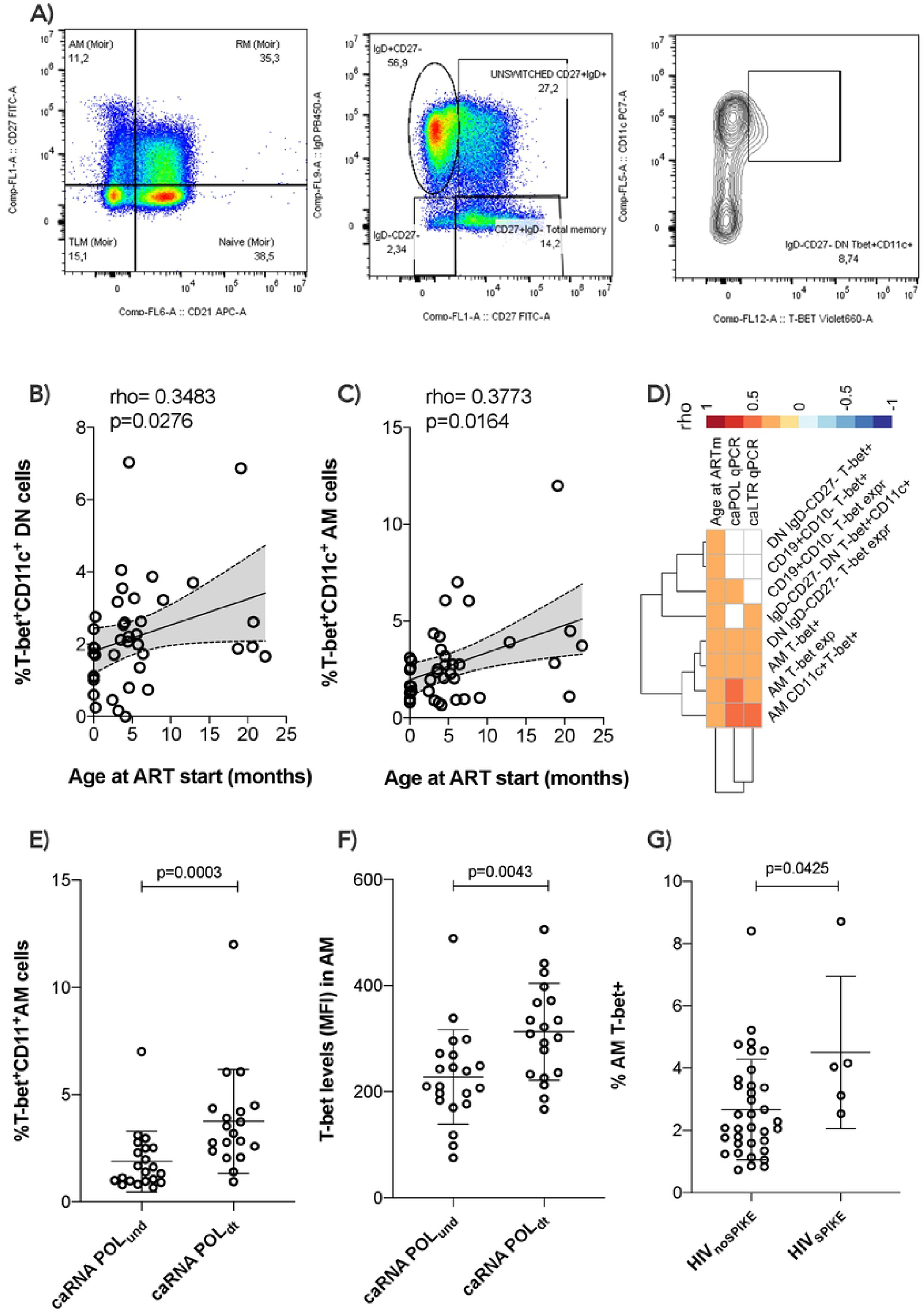
Time of ART initiation and cell associated HIV-1 RNA (caRNA) are associated with expansion of aging B cells. Gating strategy is shown in a); in b) and c) correlations between aging B cells and age at ART start are shown; d) correlation plot between viral correlates of recent replication and aging B-cells / exhausted T-cells are shown; differential analysis between levels of aging B cells and caRNA or SPIKE being detected vs non-detected is shown in e), f), g), p values are calculated using Mann Whitney test. Spearman p values are shown in b), c), and d). Significance was set at p>0.05. DN= double negative; AM= activated memory; MFI= mean fluorescent intensity.

### Time of ART-start and caRNA are associated with levels of aging T-bet+ CD11c+ B cells

We performed an extensive immune phenotyping focusing on the B-cell compartment (gating strategy shown in Fig.1A). We found that the proportion of DN and AM expressing both T-bet and CD11c were positively associated with the time of ART initiation, with expansion of T-bet^+^CD11c^+^DN B cells (p=0.03, Fig. 1B) and T-bet^+^CD11c^+^ AM B cells (p=0.02, Fig.1C) in those with delayed ART initiation. We further explored if the levels of aging in the B-cell compartment could be associated with the HIV-reservoir. Whereas caHIV-1 DNA showed no association with any evidence of aging B cell compartment, both total caHIV-1 RNA (LTR) and unspliced caHIV-1 RNA (Pol) demonstrated a positive association with the aging-B cells (Fig. 1D). caHIV-1 RNA was associated with B cells, AM and DN expressing T-bet^+^ alone or together with CD11c, with higher levels of these B-cell populations present in individuals with detectable ongoing virus expression (Fig. 1E, 1F). We further stratified the study participants by those who did (group I= 5) or did not (group II= 35) experience Spikes during their lifetime (Fig. 1g). Group I had significantly higher levels of AM T-bet+ cells compared to group II (p=0.04, Fig. 1g). These data showed that age at ART initiation is strongly correlated with levels of B-cell aging in PHIV and that ongoing HIV-1 replication is associated with precocious aging.

## Individuals with expansion of aging B cells have elevated levels of exhausted T-cells

We then explored whether the levels of aging within the B-cell compartment corresponded to elevated levels of exhausted T-cells. Within the aging B-cells we included T-bet^+^CD11c^+^, T-bet+ only B-cells, or levels of T-bet (MFI) within the whole B compartment as well as within the ‘namely aging’ phenotypes (AM and DN). In assessing the T-cell compartment, we focused on populations expressing exhaustion biomarkers (Fig. 2). Overall, correlation analysis demonstrated direct positive association between B and T cells, suggesting that a certain extent of immune aging/exhaustion persisted in different cellular populations, even many years after successful treatment and virological control. AM T-bet^+^CD11c^+^ was associated with PD-1 expression on CD4 effector (p=0.006) and T follicular helper cells (Tfh) (p=0.049) cells. Furthermore, TIGIT expression on CD4 subset and on Tfh showed a strong positive association with all the aging B-cell populations (Fig. 2B). Similarly, LAG3 expression on transitional memory (TTM) demonstrated a strong association with AM (p=0.002) and DN (p=0.003) expressing both T-bet and CD11c. These data demonstrated that premature aging and exhaustion persists simultaneously in both B and T cell compartments, even after >10 years of ART.

**Fig.2.**
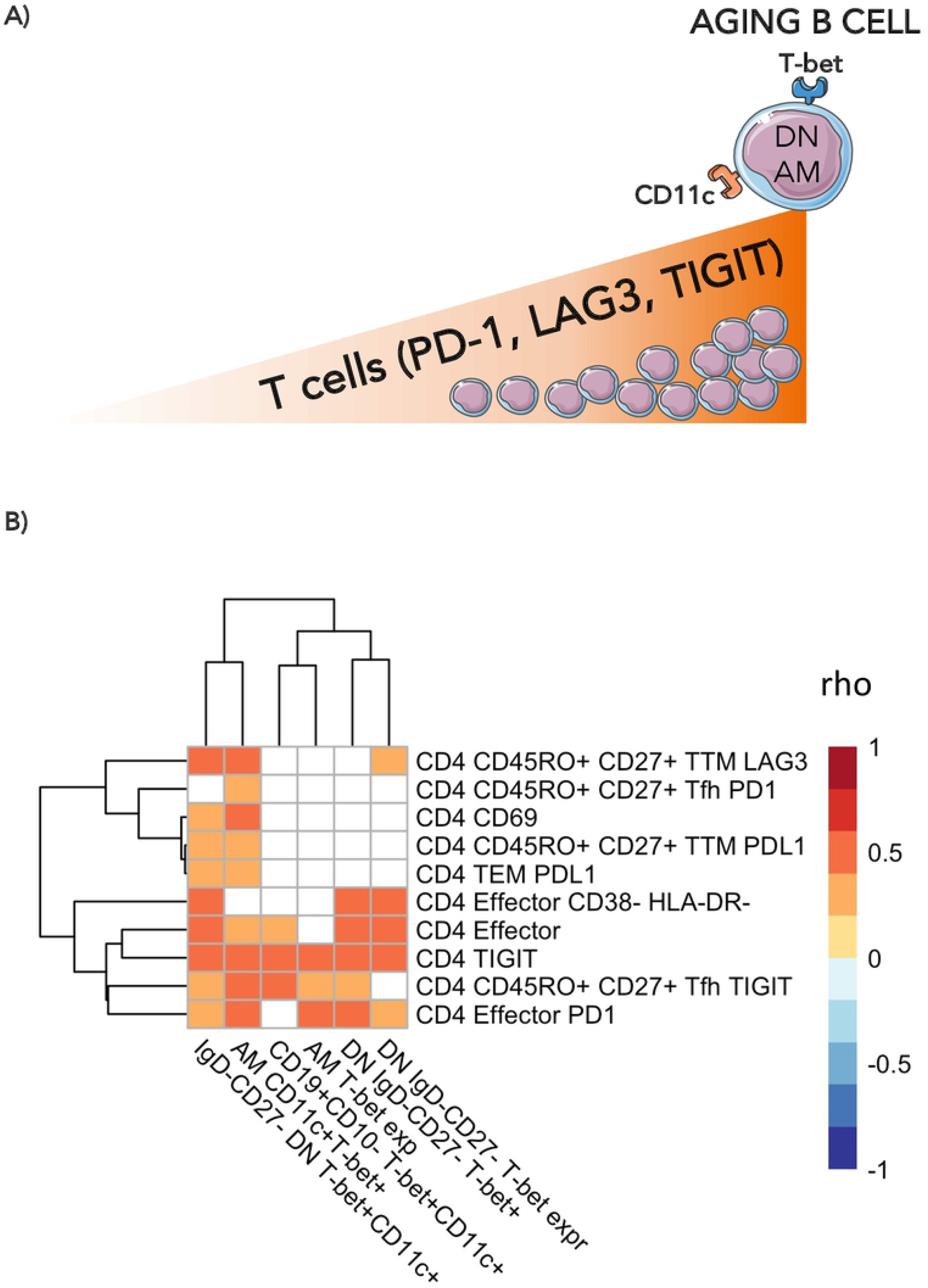
Levels of exhausted T-cells are positively associated with levels of aging B-cells. In a) a cartoon showing the main findings of the figures are pictured. In b) Heatmap plot showing Spearman correlations between exhausted T-cells and levels of aging B-cells. Only significant correlations are shown with red indicating positive correlations and Blue the negative ones. The colored scale going between 1 and -1 indicates the rho values. DN= double negative; AM= activated memory. Significance was set at p<0.05.

### Proteomic profiles associated with precocious immune aging

To assess whether humoral/soluble factors might correspond to aging/exhaustion phenotypes, we performed liquid chromatography/mass spectrometry-based proteomics, detecting 338 plasma proteins (16). The distinct immunological, virological, and clinical features associated with immune aging were correlated to the whole plasma proteomic profile (Supp. Fig 1). Two distinct clusters were initially identified which were negatively (36 proteins) or positively (37 proteins) associated with features of immune exhaustion (Fig. 3A). Such protein clusters were further interrogated for their biological role by enrichment analysis on Reactome and Gene Ontology (GO) biological processes databases (Fig. 3B). Immune inflammation and complement cascade activation pathways were enriched in proteins positively associated with features of immune aging (bottom panels, Fig. 3B and 3C). Indeed, amyloid P component in serum (APCS) and clustering (CLU), both involved in apoptotic, aging and tumor progression processes (GO:0002673) together with complement cascade molecules such as C5, CFI, C4BPA, CFB (R-HSA-173623) were positively associated to selected features of immune aging (Supplementary Table 1). In addition, proteins of light and heavy chain of immunoglobulins, involved in humoral immune response pathway (GO:0002920) such as IGLV1-47, IGHV4-34, IGLV2-23, IGHV3-48 were positively associated with immune aging. Enrichment analysis performed on negatively correlated proteins, showed no association with inflammatory pathways but only with processes involved in coagulation. Indeed, proteins such as APOH, SERPINF2, HRG involved in pathways of negative regulation of blood coagulation (GO:0030195) and platelet degranulation (R-HSA-76002) were negatively associated with features of aging (Supplementary Table 1).

**Fig 3.**
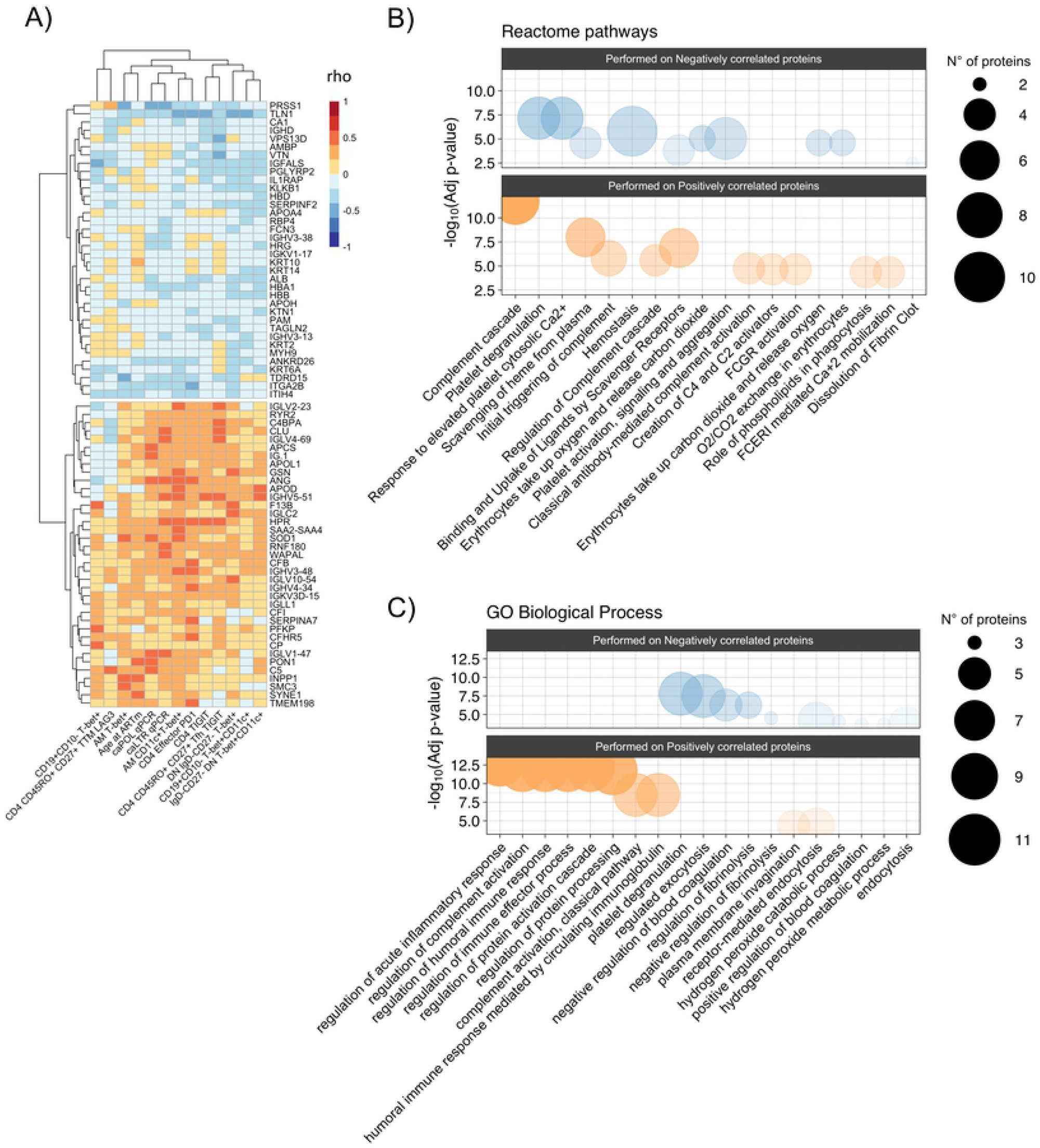
Association between proteomic profiling and levels of aging B-cells and exhausted T-cells. A) Heatmap plot showing Spearman correlations between the 13 unfunctional features values and the abondance of the 73 plasma proteins belong to the two clusters identified in correlation matrix with all 338 proteins. Red indicates positive correlations and Blue negative ones. Bubble plots showing the top 10 Reactome pathways (B) and GO Biological Process (C) significantly enriched (Adjusted p-value < 0.05) in proteins positively (Pos) and negatively (Neg) correlated with the 13 unfunctional features. The proteins were separated into positively and negatively correlated based on the two clusters showed in the correlation heatmap in panel A. Colors are related at the log_10_ adjusted p-value values and the circle diameter are related at the number of proteins for each term. Significance was set at p<0.05.

### Expansion of aging B-cells is associated with B-cell dysfunctionality in PHIV

We further assessed whether the presence of aging B-cells could affect the functionality of the B-cell compartment to maintain immunological memory against vaccinations, such as measles. Interestingly, the proportion of B-cells expressing the senescence marker T-bet, demonstrated negative association with the capacity of B-cells to maintain immunological memory to measles vaccination (Fig. 4A). Higher levels of CD19+CD10-T-bet+ B cells were associated with reduced plasma concentrations of anti-measles specific IgG (Fig. 4B, rho=-0.338, p=0.03546). Of note, his association was strong regardless of the time of ART initiation (Fig. 4C) or timing from the last booster vaccination (Fig.4D).

**Fig 4.**
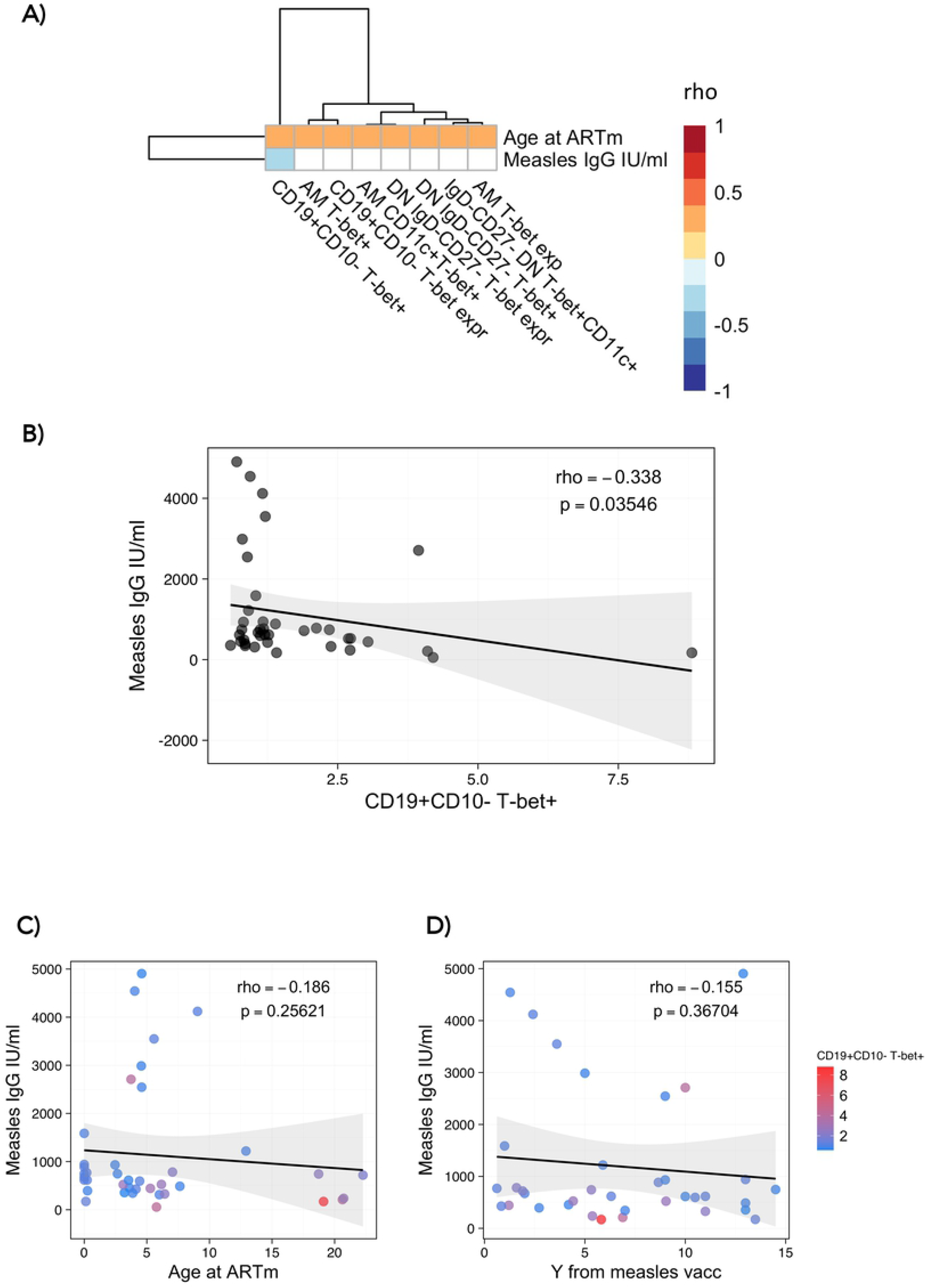
Association between aging B-cells and anti-measles humoral response. A) Heatmap plot showing Spearman correlations between aging B-cells and anti-measles plasma IgG titers (IU/ml). Red indicates positive correlations and Blue negative ones. B) Spearman correlation between CD19+CD10-B-cells T-bet+ and anti-Measle plasma IgG titers, with rho and p defining the statistical significance. C) and D) Spearman correlation between anti-Measle plasma IgG titers and Age at ART in m and years from measles vaccination, respectively, with rho and p defining the statistical significance. Color dots show the distribution of CD19+CD10-B-cells T-bet+. Significance was set at p<0.05.

## DISCUSSION

To our knowledge, ours is the first long-term follow-up study focusing on precocious B-cell aging in PHIV with long-term sustained virological suppression. We defined novel cellular and molecular factors associated with precocious aging in the B-cell compartment. We found that age at ART initiation, HIV caRNA, levels of exhausted T-cells and specific proteomic profiles demonstrated a strong and positive association with the levels of aging B-cells expressing T-bet alone or together with CD11c. The expansion of precocious aging B-cells appeared to have a direct impact on the ability of these patients to maintain vaccine induced immunity over time.

PHIV children, particularly younger ones, are immunologically distinct from adults including with respect to plasticity and immune regulation, resulting in a lower immune activation state (17). Since chronic immune activation and aging in treated HIV infection is probably driven by residual HIV replication (18, 19), it could be hypothesized that a prompt initiation of ART early in life followed by a sustained suppression of the viral replication may be able to minimize this (20). In this work, we show that perinatally infected adolescents growing with HIV present higher frequency of aging-B cells directly related to time of ART initiation, despite a history of continuous viral suppression, documented with at least four HIV-RNA PCR tests per year for over 10 years.

We next explored the virological determinants of the expansion of aging B-cell populations in those with PHIV. Total HIV-1 DNA did correlate with markers of B-cells aging, probably reflecting the fact that the contribution of the replication-competent virus is diluted within the entire integrated virus reservoir, which is mainly inactive (21). We thus further explore the markers of recent virus replication. Both spliced and unspliced HIV-1 caRNA were (AB) strongly associated with levels of aging B-cells. Spliced HIV-1 RNA may reflect abortive HIV-replication, with only a minor part being released as virus protein or exosome-associated fragments of RNA that can still trigger immune activation (22). In contrast, the unspliced HIV-1 RNA is thought to predict the replicative-competence of the virus reservoirs and has been associated with virologic failure and markers of immune activation in elite controllers (20, 23, 24), recently proposed as a predictive marker of viral rebound (25). In our cohort, aging B-cells were not only associated with caRNA, but frequency of aging B-cells was higher in those PHIV adolescents who experienced HIV spikes in absence of virologic failure. The association between expansion of aging B-cell, caRNa and viral Spikes is consistent with the hypothesis that precocious aging in the B cell compartment is dependent on HIV-1 replication and virus particle release, which fuels chronic immune activation, exhaustion and ultimately aging (26).

Multiple mechanisms likely underpin the association between caHIV-1 RNA and aging B-cells: 1) HIV-1 particles can interact directly with B cells surface-bound via the CD21 receptor with complement 3 (C3) fragment both in peripheral blood and lymph nodes of HIV-1 patients (Kardava L. et al. 2018); and 2) B-cells may function as Antigen Presenting Cells (APC) taking direct contact with follicular T-cells to trigger an anti-HIV-response. In case of HIV-persistence, both B and T cells should experience a state of chronic immune activation resulting in expansion of signatures associated with precocious aging (27, 28). Consistent with this hypothesis, our results showed that aging B-cells existed simultaneously with T-cell exhaustion. T-bet+CD11c+ B-cells showed strong association with T cells expressing PD-1, TIM-3 and LAG-3 which are inhibitory receptors that are found to be increased on the T-cell surface as a consequence of persistent activation and described as markers of cells exhaustion (29). Furthermore, T-bet+CD11c+ aging B-cells were associated with exhausted Tfh in accordance with other models of chronic antigenic stimulation such as auto-immune diseases (30). In fact, the excessive T-bet+CD11c+ age-associated B cells (ABCs) (31) not only to contribute to the production of auto-Abs but also to promote aberrant Tfh cell differentiation and consequently compromising affinity-based germinal center B-cell selection and Ab-affinity maturation in lupus mouse models.

There are very likely other modes of soluble factor-receptor interactions which can regulate B cells during HIV-1 infection and may contribute to progression to aging of B-cell compartment (32). To assess this possibility, we analyzed proteomic profiles of our patients, defining at the plasma level the status of immune activation and precocious aging found in B and T cell phenotype analysis. Proteins positively associated with features of HIV-related immune exhaustion were mainly involved in pro-inflammatory and complement activation processes. While it was previously shown that the early initiation of suppressive ART over the acute phase of the infection in HIV-infected adults reduced aspects of the immune activation (18, 19), we here show the persistence of bio humoral correlates of exhaustion and aging in PHIV with a history of long-term viral suppression (>10 years). Specifically, APCS and CLU, both involved in processes of cell apoptosis, inflammation, and lymphoproliferative processes (33-35) were positively associated to caHIV-RNA, immune checkpoint-inhibitors (TIGIT and PD1 on T cells) and exhausted B cells (T-bet+CD11c+ B cell subsets). Accordingly, such proteins were shown to be higher in virally controlled HIV infected adult experiencing a poor immune reconstitution and disease progression despite viral control (36).

Proteomics further showed that complement cascade activation pathway was enriched in proteins positively associated with immunological aging features including CLU. As previously demonstrated, the complement activation contributes to a chronic pro-inflammatory environment even in well-controlled HIV infected adults (37). Whereas the activation of the complement cascade during acute HIV infection is largely via activation of the classical pathway (36, 38), recent studies highlight how complement factors bind IgG3 on exhausted B cell subsets (TLM) in HIV-positive individuals (32, 39). In line with this evidence, our results showed a positive association of both caHIV-RNA and aging B cell subsets (T-bet+CD11c+ DN and AM) with plasma complement cascade proteins. Correlation analysis further revealed an association of proteins involved in coagulation processes with features of immune aging. As previously shown in adults, a pro-coagulative imbalance, partially resolved by ART initiation during the acute infection (18, 19) and persisting over time in HIV infected adults (40), was confirmed in our cohort where a regulation of fibrinolysis was negatively associated with features of aging in both T and B cell compartment. Overall, plasma proteomic profiling may suggest that the persistence of complement cascade perturbation, rather than inflammatory and coagulation proteins may contribute to B -cell exhaustion and signs of precocious aging in long term virally controlled (>10 years) PHIV.

Finally, we explored if the presence of this expanded aging B-cell population could reflect an impairment of the maintenance of the humoral response towards childhood vaccination, such as measles immunization which should be maintained throughout life in physiological conditions. We found that levels of T-bet on the global B-cell population were negatively associated with anti-measles serum IgG. These data are not due to the natural Ab decay because patients were analyzed at similar median years from vaccination. Our observations raise the possibility that the maintenance of specific Ab titers is related to a better maturation and preservation of the memory B cell compartment as a direct consequence of an early ART.

In conclusion, our study demonstrated for the first time the impact of a late ART start on B-cell compartment is still visible despite >10 years of suppressive ART. This set of data also suggests a role of T-bet and CD11c towards the definition of B-cell exhaustion in PHIV and showed that the subset of T-bet expressing B cells may negatively affect the capacity of B-cell compartment to maintain a vaccine-induced functional Ab response. Further studies aiming to confirm whether such multi-omic signatures of aging/inflammation can inform simplified methods to stratify risk of disease progression or lymphoproliferative disorders in cohorts of long-term suppressed PHIV are needed.

### Limitations of the study

While our study featured multiple strengths, as with all research it also had a number of important limitations, including: a) the small size of the CARMA cohort limited the power of correlation analysis to detect associations; it would be interesting to expand the immunological profiling to a larger cohort; b) the lack of a control group of exposed uninfected HIV individuals and potentially of another group that started therapy after 2 years of age, to deeply investigate the impact of late ART start; and c) the cross-sectional study design.

## Acknowledgements

We would like to acknowledge all patients and guardians who decided to participate in the study. We thank Jennifer Faudella and Giulia Neccia for her administrative assistance. We thank Nadia Iavarone and Tamara Di Marco for their nursing assistance. We thank Ilaria Pepponi for her precious lab management work.

## Author Contribution

**Conceptualization:** Paolo Palma, Alessandra Ruggiero, Giuseppe Rubens Pascucci, Nicola Cotugno.

**Data curation:** Sara Dominguez-Rodrıguez, Giuseppe Rubens Pascucci

**Formal analysis:** Alessandra Ruggiero, Giuseppe Rubens Pascucci, Sara Dominguez-Rodrıguez.

**Funding acquisition:** Carlo Giaquinto, Paolo Rossi, Paolo Palma, Nicola Cotugno

**Investigation:** Alessandra Ruggiero, Giuseppe Rubens Pascucci, Nicola Cotugno, Stefano Rinaldi, Kinga Smolen, Al Ozonoff, Caroline Foster, Alasdair Bamford, Nigel Klein, Anita DeRossi, Eleni Nastouli

**Methodology:** Paolo Palma, Alessandra Ruggiero, Giuseppe Rubens Pascucci, Nicola Cotugno **Resources:** Nicola Cotugno, Pablo Rojo, Eleni Nastouli, Nigel Klein, Caroline Foster, Anita De Rossi, Carlo Giaquinto, Paolo Rossi, Savita Pahwa, Paolo Palma.

**Supervision:** Hanno Steen, Katherine Luzuriaga, Philip Goulder, Paolo Rossi, Ofer Levy, Savita Pahwa, Paolo Palma

**Visualization:** Alessandra Ruggiero, Giuseppe Rubens Pascucci, Nicola Cotugno, Paolo Palma **Writing – original draft:** Alessandra Ruggiero, Giuseppe Rubens Pascucci, Nicola Cotugno, Paolo Palma

**Writing – review & editing:** all authors

## MATERIALS AND METHODS

### Ethics statement

This is a multi-center study which include the following institutions: Bambino Gesù Children’s Hospital (OPBG, Rome, Italy), University of Padua (Padova, Italy), University Hospital 12 de Octubre (Madrid, Spain), Hospital Gregorio Marañón (Madrid, Spain), Imperial College Healthcare NHS Trust (London, UK), Great Ormond Street Hospital (London, UK), Brighton and Sussex University Hospitals (Brighton, UK). Each recruiting sites received approval by local ethic committees (Foster, Dominguez-Rodriguez et al. 2020). Study participants or their legal guardians gave written informed consent in accordance with the Declaration of Helsinki.

### Study population

The CARMA (Child and Adolescent Reservoir Measurements on early suppressive ART) cohort is part of the existing EPIICAL consortium (Early treated Perinatally HIV Infected individuals: Improving Children’s Actual Life) (41, 42), a multi-center, multi-cohort global collaboration primarily supported by PENTA foundation (Pediatric European Network for the Treatment of AIDS). CARMA included 40 perinatally HIV infected children (PHIV) with following inclusion criteria: (1) start of ART within the 2^nd^ year of life; (2) ≥5 years of age; (3) viral suppression (<400 copies/mL) achieved in the first 12 months after initiation of ART and maintained for at least 5 years with 4 plasma viral load tests performed each year prior to enrolment; (4) A single viral load between 400 and 1000 c/mL (Spike) is permitted annually returning to less than 50 c/ml on next testing (within 3 months); (5) plasma viral load of <50 HIV-1 RNA copies/ml at enrolment. Wider characteristics of participants were described elsewhere (41) and relevant info provided in Table 1. CD4 counts were collected at the hospital visits and vaccination history was available from patients’ files.

### Samples collection

Plasma samples were obtained by centrifugation of EDTA-blood at 2000xg for 10’ and stored at - 80°C until use. Peripheral blood mononuclear cells (PBMCs) were isolated using Ficoll density gradient centrifugation, resuspended in fetal bovine serum (FBS) supplemented with 10% dimethyl sulfoxide (DMSO) and stored in liquid nitrogen until use.

### B and T-cell phenotypic analysis

PBMCs from 40 PHIV were thawed, washed, and stained with the LIVE/DEAD fixable BV510 dead cell stain kit according to manufacturer’s protocol (Life Technologies, Carlsbad, CA), used to assess viability: positive cells were thus excluded from the analysis as they were considered as dead. For B-cell phenotype, after washing with PBS 10% FBS, cells underwent surface staining with the following monoclonal antibodies (mAbs, from BD Biosciences): CD3, CD10, CD16 (BV510), CD19 (APC-R700), CD21 (APC), CD27 (FITC), IgD (BV421), IgM (PE-CF594), IgG (BV605), CD11C (PC-7).

Finally, stained cells were resuspended in 1% paraformaldehyde (PFA) and acquired using Stained cells were acquired on Cytoflex (Beckman Coulter, Brea, CA) and analysed with FlowJo v10.0.8 (Tree Star) software. Following surface staining fixing and permeabilization of cells (BD permeabilization solution II 1x), cells were stained with an anti T-bet BV650 (04-46, BD). For T-cell phenotype, LIVE/DEAD Fixable Blue Dead Cell Stain Kit from Thermo Fisher Scientific (Boston, MA) was used to detect and exclude dead cells. After washing with PBS 10% FBS, cells underwent surface staining with the following monoclonal antibodies as previously described (Rinaldi S. et al. 2021): LAG3 BV650, TIGIT PE-Cy7, CD19 Alexa Fluor 700, HLA-DR PE, CCR7 FITC, CD38 BV711, PD-L1 BV711, PD-1 BV421, and CD8 PerCP from BioLegend (San Diego, CA); CD3 BUV496, CD4 APC-Cy7, CD4 APC-H7, PD-1 BV650, CXCR5 Alexa Fluor 647, and CD27 BV480 from BD Biosciences (San Jose, CA); and CD45RO PE-Cy5.5 from Beckman Coulter (Fullerton, CA). Finally, stained cells were resuspended in 1% paraformaldehyde (PFA) and acquired using Stained cells were acquired on a BD LSRFortessa (BD Biosciences) and analysis performed using FlowJo v10.0.8 (Tree Star) software. Gating strategies for B-cell phenotypes, T-bet and CD11c are provided in Fig.1. Gating strategies for T-cell analysis were shown previously (28). Positive cell gating was set using fluorescence minus one control. All the reagents were tested and titrated for optimum concentration before usage.

### Quantitative total HIV-1 DNA assay

Total HIV-1 DNA was quantified in PBMCs of 40 PHIV by real-time quantitative reverse transcription PCR (qRT-PCR) as previously described (43). All measurements were done in triplicates. Results are reported as copies of HIV-1 per million cells.

### Quantitative caHIV-1 RNA assay

caHIV-1 RNA was quantified as described in (28). Briefly, Qiasymphony automated platform was used to isolate total cellular RNA (DSP virus/pathogen mini kit (Qiagen). RNA was further processed in an in-house assay using primers of previously validated assays (44, 45) to selectively amplify total (LTR) and unspliced (pol) ca-HIV-1 RNA via qRT-PCR. In order to express caHIV-1 RNA copies per 10^6^ PBMC, the caHIV-1 RNA measurements were normalized against cellular genes TBP1 and IPO8 expression.

### Plasma proteomics preparation and analysis

Plasma proteomics data was produced using a High-performance liquid chromatography mass spectrometry (HPLC/MS) method as previously described (16). The sample processing employed an MStern blotting protocol previously developed and validated *in house* (46-49). In brief, 1 μL of plasma (∼50 μg of proteins) was mixed in 100 μL of urea buffer. Following reduction and alkylation of the cysteine side chains, an amount of 15 μg of proteins was loaded on to a 96-well plate with a polyvinylidene fluoride (PVDF) membrane at the bottom (Millipore-Sigma), which had been previously activated and primed. Trypsinization of the proteins adsorbed to the membrane was achieved by incubation with the protease for 2h at 37°C. Resulting tryptic peptides were eluted off the membrane with 40% acetonitrile (ACN)/0.1% formic acid (FA). The peptides were subsequently cleaned-up using a 96-well MACROSPIN C18 plate (TARGA, The NestGroup Inc.). The samples were analysed on the same LC/MS system as the data-dependent acquisition (DDA) runs using identical LC parameters (45 minutes gradient, 59 minutes total runtime). The m/z range 375−1200, covering 95% of the identified peptide, was divided into 15 variable windows based on density, and the following parameters were used for the subsequent DIA analysis: resolution 35000 @ m/z 200, AGC target 3e6, maximum IT 120 ms, fixed first mass m/z 200, NCE 27. The DIA scans preceded an MS1 Full scan with identical parameters yielding a total cycle time of 2.4s. We use a previously published in house generated spectral library (16). All DIA data were directly analysed in Spectronaut v12.0.20491.18 (Biognosys, Switzerland). Standard search settings were employed, which included enabling dynamic peak detection, automatic precision nonlinear iRT calibration, interference correction, and cross run normalization (total peak area). All results were filtered by a q-value of 0.01 (corresponding to an FDR of 1% on the precursor and protein levels). Otherwise default settings were used.

### Anti-Measles IgG

Plasma Anti-Measles IgG titres were measures using EuroImmunAnti-Morbillo ELISA (IgG) (LOT E180111AE), following manufactures instruction. Results given as UI/L.

### Statistical analyses

Between-group comparisons were performed using non-parametric U-Mann-Whitney test for continuous variables or Fisher’s exact test for categorical variables. Spearman correlation (rho) was used to describe the association between continuous variables. Proteins and cell populations with >70% zero values or >50% missing data were omitted from heatmaps. To focus on single associations (Fig. 1d, 2b and 3a) only statistically significant correlations (p-values <0.05) were shown. In other cases, to highlight clustering patterns, were shown all correlations (Fig. 3a and Supp Fig. 1). The chromatic scale is proportional to the Spearman correlation, using red for positive correlations (rho > 0) and blue for negative ones (rho < 0). To investigate the biological role of the proteins belonging to the two clusters (Fig 3a), a pathway enrichment analysis in Reactome 2016 and GO Biological Process 2021 databases was performed using the R package “enrichr” v3.0 (50). Statistical analyses were performed using R (version 4.1.1) or GraphPad Prism 6.0 software (San Diego, CA).

## Supporting information captions

**Supplementary Fig. 1 Heatmaps showing Spearman correlations between abundance of 338 plasma proteins and 13 selected unfunctional features**. Red indicates positive correlations and Blue negative ones.

**Supplementary Table. 1 Overview of proteins within the Reactome and GO biological process pathways associated with aging B-cells and exhausted T-cells**.

